# A small dynamic leaf-level model predicting photosynthesis in greenhouse tomatoes

**DOI:** 10.1101/2022.09.10.507401

**Authors:** D Joubert, N Zhang, S.R. Berman, E Kaiser, J Molenaar, J.D. Stigter

**Affiliations:** Mathematical and Statistical Methods Group, Wageningen University and Research, Wageningen, The Netherlands; Horticulture and Product Physiology, Wageningen University and Research, Wageningen, The Netherlands

## Abstract

The conversion of supplemental greenhouse light energy into biomass is not always optimal. Recent trends in global energy prices and discussions on climate change highlight the need to reduce our energy footprint associated with the use of supplemental light in greenhouse crop production. This can be achieved by implementing “smart” lighting regimens which in turn rely on a good understanding of how fluctuating light influences photosynthetic physiology.

Here, a simple fit-for-purpose dynamic model is presented. It accurately predicts net leaf photosynthesis under natural fluctuating light. It comprises two ordinary differential equations predicting: 1) the total stomatal conductance to CO_2_ diffusion and 2) the CO_2_ concentration inside a leaf. It contains elements of the Farquhar-von Caemmerer-Berry model and the successful incorporation of this model suggests that for tomato (*Solanum lycopersicum L*.), it is sufficient to assume that Rubisco remains activated despite rapid fluctuations in irradiance. Furthermore, predictions of the net photosynthetic rate under both 400ppm and enriched 800ppm ambient CO_2_ concentrations indicate a strong correlation between the dynamic rate of photosynthesis and the rate of electron transport. Finally, we are able to indicate whether dynamic photosynthesis is Rubisco or electron transport rate limited.

**Author summary:** The cultivation of greenhouse crops under optimised conditions will become increasingly important, with supplemental lighting playing a vital role. However, converting light energy into plant photosynthesis is not always optimal. A potential venue that may lead to the efficient conversion of light energy involves a model-based implementation of “smart” lighting control strategy. This approach does however necessitate a good understanding of how plants harness light energy under natural fluctuating irradiance. Accordingly, as a first step, we have developed a small leaf-level model that predicts dynamic photosynthesis in natural fluctuating light. It may potentially be used in future supplemental light control applications.

## Introduction

The cultivation of greenhouse crops under optimised conditions will become increasingly important, with the need for year-round crop harvesting under changing environmental conditions as a driving factor. The widespread use of supplemental lighting to optimise growing conditions is a key tool at our disposal. The efficiency of converting light energy into plant photosynthesis is however not always optimal. This is particularly true when lights are first turned on, where time is required for the activation of key photosynthetic enzymes and for adjustments in stomatal pore aperture [1].

Given recent trends in global energy prices and continuous discussions on climate change, our energy footprint associated with the use of supplemental light needs to be reduced. Two potential avenues that may lead to the efficient conversion of light energy into photosynthesis are: 1) the use of light-emitting diodes (LEDs) in supplemental lighting applications. LEDs increase the efficiency with which electrical energy is converted to photosynthetically active radiation (PAR), the radiation with wavelength 400-700nm, which powers photosynthesis [2]. 2) A model-based implementation of “smart” lighting regimens. This approach necessitates a good understanding of how plants harness light energy under natural fluctuating irradiance (I).

Plant responses to fluctuating irradiance occur across numerous levels of complexity, ranging from the whole canopy (at crop level and governed by plant structure) to the biochemistry of a single reaction (at leaf level and governed by physiology) and across different orders of magnitude in time. At the top of the canopy light levels depend on factors such as the solar angle and amount of cloud cover. Lower parts of a crop canopy rely on light in the form of sun-flecks, and the amount of these in turn depends on the canopy’s structure [3].

Light intensity is the most dynamic condition to which greenhouse crops need to respond and can change at time scales ranging from a season (winter versus summer) to less than 1 second (passing clouds) [4]. The variation of incident light on leaves in the upper canopy can have a substantial effect on photosynthesis in this upper layer since it accounts for up to 75% of crop canopy carbon assimilation [5].

At leaf level, a plant’s ability to regulate photosynthesis in response to rapid variations in irradiance may be restricted by the following factors: 1) *the opening/closing of stomata*, 2) *the activation/deactivation of Calvin cycle enzymes*, 3) *the up-regulation/down-regulation of photoprotective processes* [3], and 4) *transiently changing mesophyllic conductance* [6].

Previous leaf-level models have ranged both in complexity and in time scales of prediction. These often include the well-know Farquhar-von Caemmerer-Berry (FvCB) model, which mathematically describes key Calvin cycle processes and linear transport rates [7]. The result is an estimated net photosynthetic rate (A_n_) which stems from competitive enzymatic processes involving CO_2_ and O_2_ binding under different environmental conditions [8]. A brief summary of selected *small* dynamic models is given in Table (1). We included a summary of the model presented here for comparison.

**Table 1.**
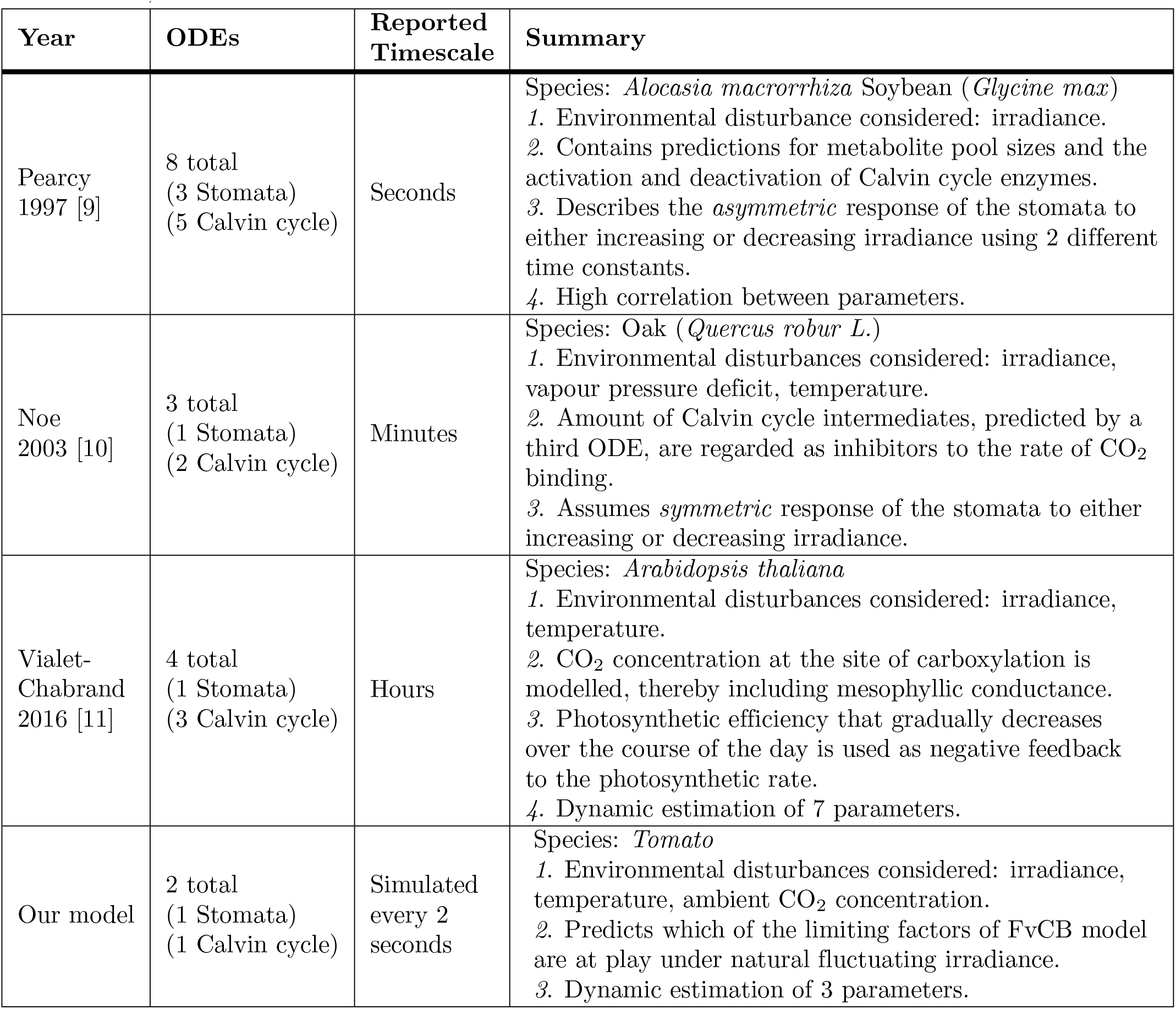
Summary of selected small dynamic leaf-level photosynthesis models. (Ordinary differential equations abbreviated as ODEs).

| Year | ODEs | Reported Timescale | Summary |
| --- | --- | --- | --- |
| Pearcy<br>1997 [9] | 8 total<br>(3 Stomata)<br>(5 Calvin cycle) | Seconds | Species: *Alocasia macrorrhiza* Soybean (*Glycine max*)<br>1. Environmental disturbance considered: irradiance.<br>2. Contains predictions for metabolite pool sizes and the activation and deactivation of Calvin cycle enzymes.<br>3. Describes the *asymmetric* response of the stomata to either increasing or decreasing irradiance using 2 different time constants.<br>4. High correlation between parameters. |
| Noe<br>2003 [10] | 3 total<br>(1 Stomata)<br>(2 Calvin cycle) | Minutes | Species: Oak (*Quercus robur L.*)<br>1. Environmental disturbances considered: irradiance, vapour pressure deficit, temperature.<br>2. Amount of Calvin cycle intermediates, predicted by a third ODE, are regarded as inhibitors to the rate of CO<sub>2</sub> binding.<br>3. Assumes *symmetric* response of the stomata to either increasing or decreasing irradiance. |
| Vialet-Chabrand<br>2016 [11] | 4 total<br>(1 Stomata)<br>(3 Calvin cycle) | Hours | Species: *Arabidopsis thaliana*<br>1. Environmental disturbances considered: irradiance, temperature.<br>2. CO<sub>2</sub> concentration at the site of carboxylation is modelled, thereby including mesophyll conductance.<br>3. Photosynthetic efficiency that gradually decreases over the course of the day is used as negative feedback to the photosynthetic rate.<br>4. Dynamic estimation of 7 parameters. |
| Our model | 2 total<br>(1 Stomata)<br>(1 Calvin cycle) | Simulated every 2 seconds | Species: *Tomato*<br>1. Environmental disturbances considered: irradiance, temperature, ambient CO<sub>2</sub> concentration.<br>2. Predicts which of the limiting factors of FvCB model are at play under natural fluctuating irradiance.<br>3. Dynamic estimation of 3 parameters. |

The rest of this article is organised as follows. In section 1 we provide the theoretical background to the model and describe the plant physiology underpinning the well-known FvCB model. We also discuss three of the factors that may restrict A_n_ in greater detail. The model is defined in section 2, and the materials and methods used are discussed in section 3. Results and discussion are presented in sections 4 and 5 respectively, and conclusions are given in section 6.

## 1 Theory

We briefly introduce three of the factors that affect A_n_, and which are included in the model: 1) stomatal conductance, 2) the Rubisco limited carboxylation rate, and 3) the electron transport limited carboxylation rate.

### 1.1 The opening and closing of stomata

Stomata are located on both the upper (u) and lower (l) surfaces of tomato leaves. They are tasked with regulating the flux of gaseous H_2_O and CO_2_ between a leaf and its surroundings. They do so by adjusting their pore aperture and this is achieved by changing the form of their 2 guard cells. These structures can therefore be thought of as conductors of CO_2_ diffusion, and therefore only allow a certain amount of CO_2_ to enter a leaf.

Stomatal aperture depends on numerous environmental cues. For C3 plants (plants that allow for the direct carbon fixation of CO_2_) under non-limiting growth conditions, pore sizes increase with increased *irradiance* and low CO_2_ concentrations [12].

The metabolic regulation of these guard cells is highly complex and accordingly, an empirical model that predicts dynamic changes in the conductance of CO_2_ related to perturbations in irradiance is used here [11]. Refer to S1 Supporting Information for a discussion on how the total stomatal conductance to CO_2_ diffusion (g_tc_) is defined.

### 1.2 The FvCB model

This model describes the steady-state net photosynthetic (A_n_) rate as [7, 13]:

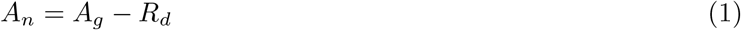

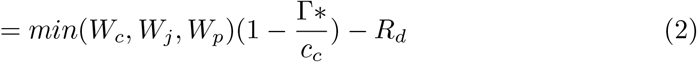

where (1) defines A_n_ as the difference between the gross photosynthetic rate (A_g_) and R_d_, the mitochondrial respiration which includes the release of CO_2_ in light other than photo-respiration [13]. From (2) follows that A_n_ can be limited by the Rubisco limited carboxylation rate (W_c_), the electron transport limited carboxylation rate (W_j_) or the rate at which triose phosphates are utilised (W_p_). Opting to keep our model structure concise and the number of unknown system parameters to a minimum, we omit the limiting factor W_p_.

#### 1.2.1 The Rubisco limited carboxylation rate (W_c_)

Once CO_2_ enters a leaf through the stomata, it diffuses through the inter-cellular spaces into the chloroplasts by means of a diffusion gradient between the chloroplast and the rest of the leaf. The numerous photosynthesis reactions that occur inside the chloroplast are summarised in the Calvin cycle, a process which comprises both light dependent and independent reactions.

During the first phase of this cycle, a single CO_2_ molecule is fixed onto a Ribulose 1,5-bisphosphate (RuBP) molecule to form *two* 3-Phosphoglyceric acid (3-PGA) molecules. Important here in the context of modelling enzyme kinetics is that this process is catalysed by the enzyme Ribulose-1,5-bisphosphate carboxylase/oxygenase (Rubisco), and its activation state is in turn increased by Rubisco activase. The structure of 3-PGA allows it to be combined and rearranged to form sugars which can be transported or stored for energy. The rate at which CO_2_ fixation takes place is known as the *carboxylation rate*. However, Rubisco also catalyses RuBP oxygenation (binding RuBP to O_2_). This reduces the efficiency of the Calvin cycle. The rate at which this takes place is called the oxygenation rate.

Mathematically, the Rubisco limited carboxylation rate is given as [7, 13, 14]:

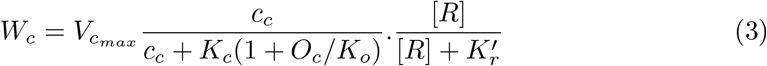

where V_cmax_ is the maximum obtainable carboxylation rate, and c_c_ is the partial pressure of CO_2_ in the chloroplast stroma whereas O_c_ is the partial pressure of O_2_ in the chloroplast stroma. K_c_ is the Michealis-Menten constant for CO_2_, K_o_ is the Michaelis-Menten constant for O_2_, [R] is the concentration of unbound (available) RuBP, and K_r_’ is the effective Michealis-Menten constant for RuBP. Assuming that RuBP is in excess, (3) reduces to [15],

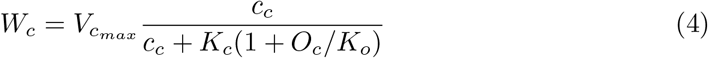

A relationship between W_c_ and the oxygenation rate is introduced in (2) by G^*∗*^, the CO_2_ concentration at which oxygenation proceeds at twice the rate of carboxylation causing the photosynthetic uptake of CO_2_ to be compensated for by the photorespiratory release of CO_2_ [16].

Notice that expression (4) is defined for CO_2_ concentrations in the chloroplast stroma (c_c_). This concentration can however not be measured directly and so is predicted if A_n_, c_i_, and g_m_ are known, by,

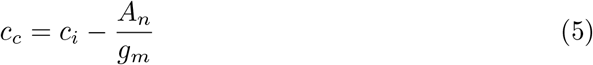

where g_*m*_ is the mesophyllic conductance encountered along the CO_2_ diffusion pathway. By assuming that this conductance is infinitely large (5) simplifies to c_c_ = c_i_.

#### 1.2.2 Electron transport limited carboxylation rate (W_j_)

The synthesis of RuBP also requires energy in the form of ATP and NADPH, and both ADP and NADP^+^ are continuously converted to these energy supplying molecules in *light dependent* reactions which are dependent on the rate of electron transport (J). Accordingly, the electron transport limited carboxylation rate (W_j_) is given as [7, 13],

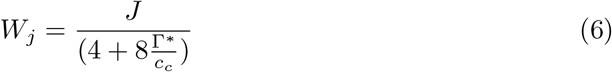

Equation (6) assumes 4 electrons per carboxylation and oxygenation and so, based on the number of electrons required for NADP+ reduction, the standard values used are 4 and 8. However, there are uncertainties in the relationship between electron transport and ATP synthesis. For example, 4.5 and 10.5 have also been used [16].

Given the above mentioned assumptions, expression (2) can be written as

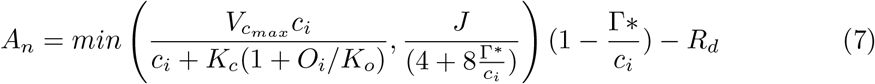

### 1.3 Temperature and light intensity effects on steady-state photosynthesis

The dependence of key FvCB model parameters on both leaf temperature (T_l_) and irradiance (I) was already included in the 1980 publication by Farquhar *et. al*. [7].

Changes in the values of parameters G^*∗*^, K_o_, and K_c_ (in (7)) associated with leaf temperature changes are described using the Arrhenius equation [16],

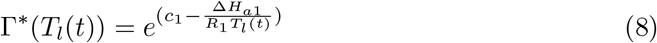

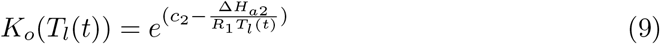

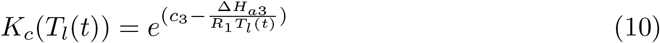

Here, the ideal gas constant R_1_, is expressed in units [kJmol^−1^K^−1^]. Constants Δ*H*_*ai*_ and c_i_, where *i* = 1, …, 3, are the respective energies of activation and scaling constants for parameters G^*∗*^, K_o_ and K_c_.

Electron transport becomes limited when insufficient quanta are absorbed [7]. Accordingly, J is modelled as a function of irradiance [17],

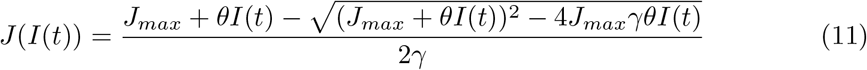

where parameter J_max_ is the upper limit to potential chloroplast electron transport determined by the components of the chloroplast electron transport chain [13].

Parameters *θ* and *γ* are unit-less (refer to S1 Supporting Information for details). All parameter values are given in Table (3).

## 2 Dynamic model structure

The model we present in this paper comprises only two ordinary differential equations, the first predicting the total stomatal conductance to CO_2_ diffusion (g_tc_) and the second the CO_2_ concentration inside the leaf (c_i_). Here, g_tc_ is the sum of the boundary layer and stomatal conductances (see S1 Supporting Information). An algebraic relation between the predicted states is used to approximate the net photosynthetic rate (A_n_). An overview of the dynamic states, system parameters, model inputs, and the measured output is given by the standard state-space representation,

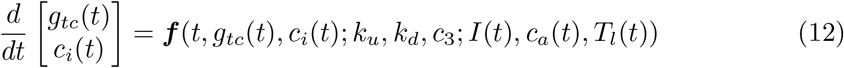

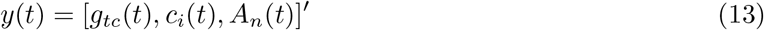

Function ***f*** is defined in equations (15)-(18). States g_tc_ and c_i_, and the predicted A_n_ are measured model outputs. The three system parameters that need to be inferred from the measured data are k_u_, k_d_ and c_3_ in expressions (10), and (15) and (16) respectively. Three environmental conditions, I(t), c_a_(t) and T_l_(t), are directly measurable and modelled as inputs/disturbances to the system. Predictions for A_n_ are made using Fick’s law of diffusion. Also known as the net flux of CO_2_ that enters a leaf, the dynamic A_n_ (achieved at a specific light intensity, as opposed to the attainable steady-state values predicted in (7)) is calculated as [18, 19]

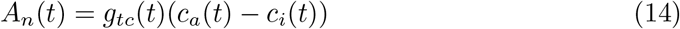

The asymmetric exponential response of stomata to increases and decreases in irradiance has often been reported [20]. This is modelled by introducing 2 time constants to the system, k_u_ describing the rate of increase in g_tc_ observed with an increase in irradiance, and k_d_ describing the rate of decrease in g_tc_ following a decrease in irradiance [9, 12, 20–24]. The resulting model structure is,

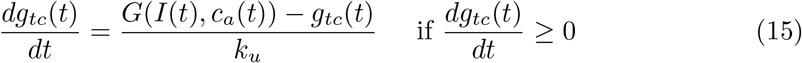

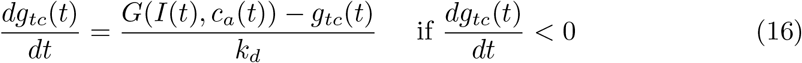

G(I(t),c_a_(t)) can be interpreted as the steady-state target function of g_tc_ for a particular combination of I(t) and c_a_(t). A description of how G(I(t),c_c_(t)) should be calibrated is given in S1 Supporting Information.

The CO_2_ concentration inside the leaf is modelled using a mass balance equation (refer to S1 Supporting Information for a discussion),

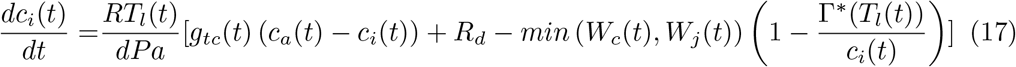

The minimum function stems from the FvCB model in equation (7). Substituting the functions W_c_(t) and W_j_(t), that have been adjusted to take T_l_ and I into account, into (17) gives,

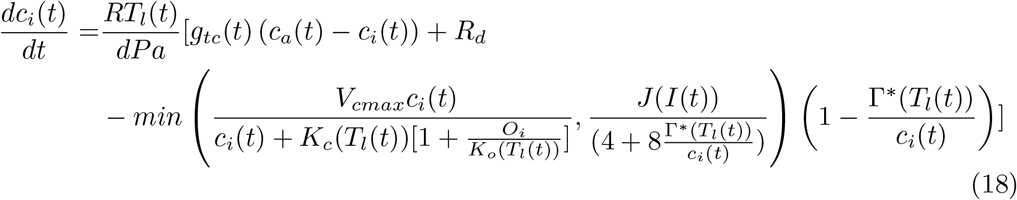

## 3 Materials and Methods

### 3.1 Growing conditions of plants

Tomtato plants were cultivated in a climate chamber (size: 16 [m^2^]) in Wageningen University & Research, Wageningen, the Netherlands (52°N, 6°E). Seeds were germinated in rockwool plugs (diameter: 2 [cm]) and transferred to rockwool cubes (10*×*10*×*7 [cm]) one week after sowing.

Unless stated otherwise, day and night temperatures were set at 23°C and 20°C respectively. Relative humidity was set at 70%. The CO_2_ concentration was kept at ambient. Plants were irrigated automatically twice per day using an ebb & flow system (at 7 AM and 7 PM) with tomato nutrient solution (EC:2.2±0.1 [*mScm*^−1^], pH:5.5) (see S1 Supporting Information).

#### 3.1.1 Plants used to estimate V_cmax_ and R_d_

These plants were exposed to an irradiance of 250 [*μmolm*^−2^*s*^−1^] provided by two types of high-pressure sodium lamps (SON-T and HPI-T PLUS, Philips Lighting). The photoperiod in the chamber was 16 hours. The SON-T lamps were switched on one hour before the HPI-T PLUS lamps and were switched off one hour after the HPI-T PLUS lamps in an attempt to mimic the gradual increase and decrease of irradiance during sunrise and sunset.

#### 3.1.2 Plants used to parameterise function J(I(t))

The plants were exposed to an average irradiance of 200 [*μmolm*^−2^*s*^−1^], with irradiance fluctuating between 50 and 500 [*μmolm*^−2^*s*^−1^] every minute. The photoperiod in the chamber was 16 hours. The irradiance pattern was randomly changed on a daily basis to simulate natural fluctuations in irradiance. Key properties including the photoperiod, minimum and maximum irradiance, daily average irradiance, and the overall shape of the light pattern were kept the same. Dynamic irradiance was provided by GreenPower LED top lighting compact modules (Philips Lighting).

#### 3.1.3 Plants used to measure photosynthesis under natural fluctuating irradiance

The plants were brought to the greenhouse compartment to grow for another four weeks. Plants were grown on growth tables in the compartment of a Venlo-type glasshouse. One layer of cloth was put on the growth table and the greenhouse compartment had a photoperiod of 16 hours to allow for ample root growth. Only when global radiation outside the greenhouse dropped below 150 [*Wm*^−2^], were high-pressure sodium (HPS) lamps (600 [W], Philips) used during the light period. These were switched off when outside global radiation increased to values above 250 [*Wm*^−2^]. When the HPS lamps were on, the light intensity from these was approximately 150 [*μmolm*^−2^*s*^−1^] at plant level. The shading screen (HARMONY 4215 O FR, Ludvig Svensson) was closed when outside global radiation increased to values above 600 [*Wm*^−2^] and was opened when outside global radiation dropped below 500 [*Wm*^−2^]. Set points of day and night temperature were 22°C and 18°C respectively. Relative humidity was set at 65% and plants were irrigated four times per day with tomato nutrient solution (see S1 Supporting Information).

### 3.2 Measurements conducted

Unless stated otherwise, gas exchange measurements were conducted on the fourth or fifth leaf of four-week-old plants (after transplanting) using a portable gas exchange system (LI-6400XT, Li-Cor Bioscience) equipped with a 6 [cm^2^] leaf chamber fluorometer. Airflow was set to 500 [*μmols*^−1^] during measurements and relative humidity was controlled at 75%. Irradiance was provided by a mixture of red (90%) and blue (10%) LEDs in the fluorometer.

#### 3.2.1 Measurements to estimate V_cmax_ and R_d_

Leaf temperature was kept around 25°C. CO_2_ response curves of photosynthesis (A_n_/c_i_ curves) were measured by changing the atmospheric CO_2_ concentration in the following order: 400, 300, 200, 100, 50, 400, 400, 500, 600, 800, 1000, 1200 ppm while keeping the light intensity at 1800 [*μmolm*^−2^*s*^−1^]. Each CO_2_ concentration step took about 2-5 minutes to finish. Measured photosynthetic rates during A_n_/c_i_ curve constructions were corrected for diffusion leaks according to the Li-Cor manual [25]. In total, eight replicates were obtained (see S1 Supporting Information for details).

#### 3.2.2 Measurements to parameterise function J(I(t))

Measurements were performed at an air temperature of 23°C, and at two atmospheric CO_2_ concentrations: 400 ppm and 800 ppm. For each CO_2_ concentration, the leaf was exposed to a respective irradiance of 0, 50, 100, 200, 400, 600, 800, 1000 and 1200 [*μmolm*^−2^*s*^−1^] for at least 45 minutes to allow both leaf net photosynthetic rate and stomatal conductance to reach steady-state. Six replicates were obtained for each CO_2_ concentration (refer to S1 Supporting Information for details).

#### 3.2.3 Measurements to track photosynthesis under natural fluctuating irradiance

Dynamic photosynthesis measurements were conducted between 3 and 24 September 2021 in a compartment (8*×*8 [m]) of a Venlo-type glasshouse located in Wageningen.

Measurements were conducted on leaves at the top of the plant that were fully exposed to sunlight and an air temperature of 23°C. Two sets of atmospheric CO_2_ concentrations, i.e. 400 ppm and 800 ppm were used. The photosynthetically active radiation (PAR) in the leaf chamber fluorometer was continuously adjusted to match the readings from an adjacent PAR sensor placed next to the chamber.

Gas exchange data were logged every two seconds between approximately 9:00 to 16:00 hours. In total, five measured replicates were obtained at 400 ppm CO_2_ and one measurement was taken at 800 ppm CO_2_.

### 3.3 Photosynthesis under naturally fluctuating irradiance: parameter estimation data set

Consider the set of observed greenhouse conditions shown in Fig.(1). The dynamics of the three model inputs, I(t), c_a_(t) and T_l_(t), observed over a 6 hour period [09:37-15:33] were measured on 8 September 2021, with c_a_(t) kept constant at 400 [*μmolm*^−2^*s*^−1^]. Notice that irradiance (Fig.(1).a) peaks during 3 stages, 30-70 [min], 105-170 [min], and again after 335 [min]. It reaches a maximum of approximately 1193 [*μmolm*^−2^*s*^−1^] for a brief period. For the majority of the day light levels fluctuate between 200 and 400 [*μmolm*^−2^*s*^−1^].

**Fig 1.**
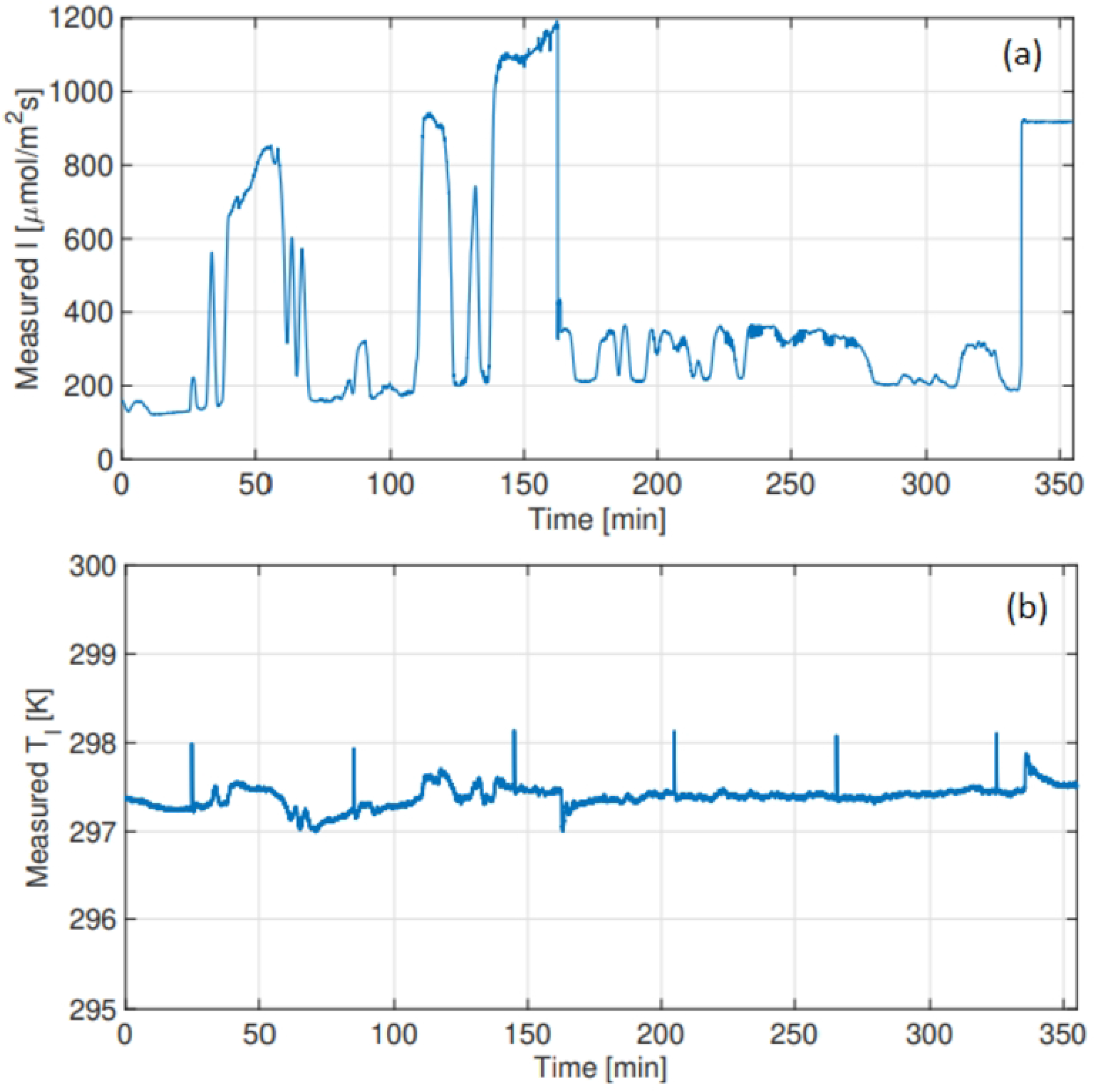
Parameter estimation data set: model inputs measured on 8 September 2021. (a) Measured irradiance (I) [*μmolm*^−2^*s*^−1^] (b) Measured leaf temperature (T_*l*_) [K].

### 3.4 Parameter Estimation

Values for the 3 unknown system parameters, k_u_, k_d_ and c_3_, were estimated using Matlab’s global optimisation function, the genetic algorithm (ga). This method is well suited to the optimisation of highly nonlinear problems and problems with a discontinuous objective function [26]. The parameters were inferred by minimising the objective function,

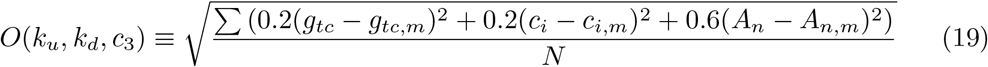

where g_tc,m_, c_i,m_ and A_n,m_ denote the measured outputs defined in (13) and N is the number of observed data points, recorded every 2 seconds. By attributing weights to the individual terms, we ensured that an accurate prediction of the important metric A_n_ was favoured. Measured g_tc_ and A_n_ values are shown in blue in Fig.(5.a) and Fig.(5.b).

*Initial conditions:* Given that both g_tc_ and c_i_ are measured outputs, the initial conditions of the 2 state equations are known.

## 4 Results

### 4.1 Estimated parameter values

Figs.(2)-(4) show the converged results of the genetic algorithm optimisation. Here, this global search method generated 300 random parameter combinations from the 3-dimensional space bounded by the intervals as indicated in Figs.(2.a)-(4.a). These results reveal the frequency with which a particular parameter value was computed as optimal. In Fig.(3.a) for example, the algorithm converged after a certain number of iterations, and computed roughly 200 of the 300 optimal points on the interval [800,850] for parameter k_d_.

**Fig 2.**
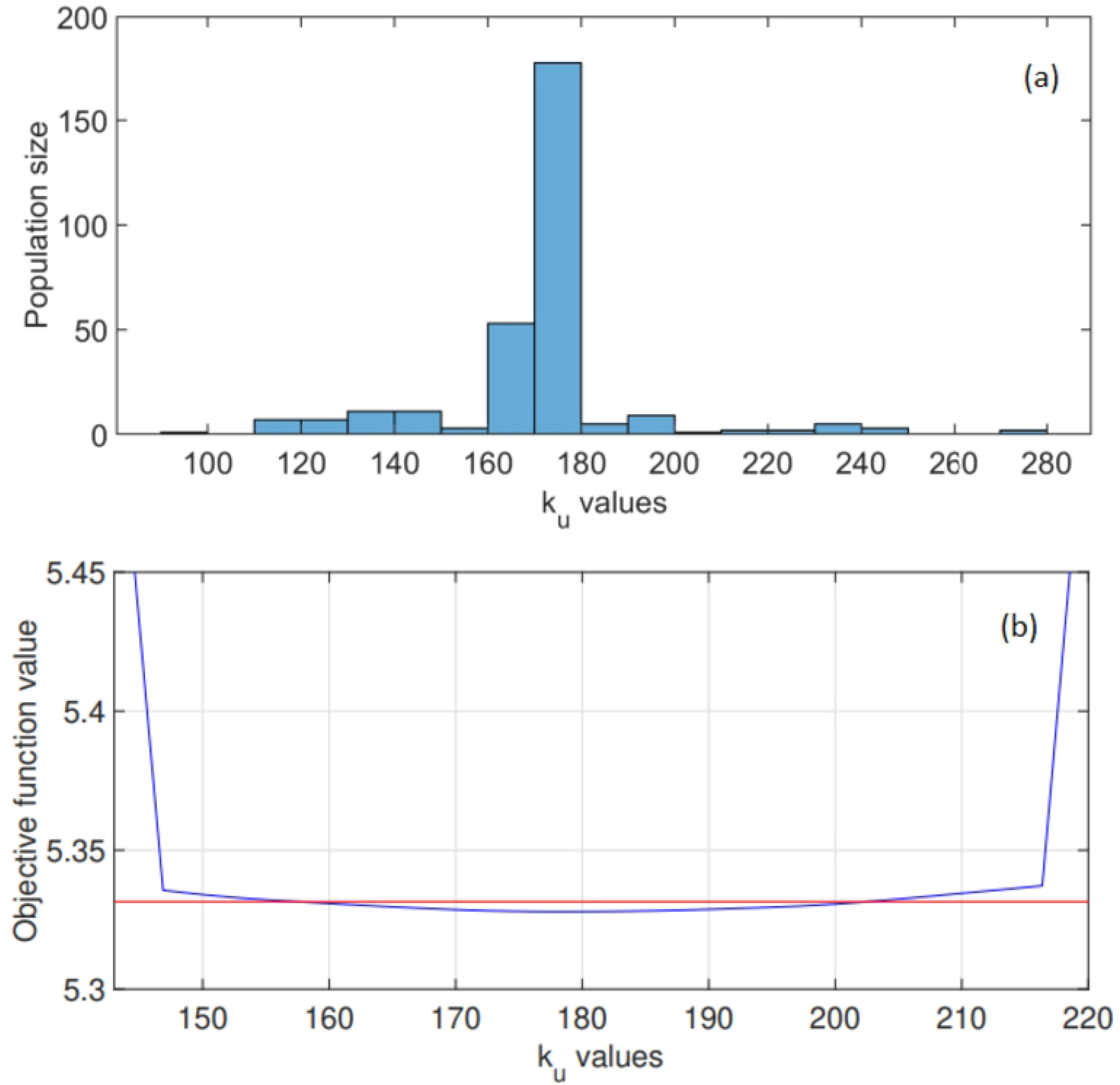
Global optimisation results for the stomatal parameter k_u_ in expression (15). (a) The range of potential k_u_ values considered is shown along with the optimised k_u_ distribution. The genetic algorithm converged to a parameter value on the interval [170,180], with the optimised k_u_=179.4 [s]. (b) Objective function values: computed profile likelihood for a range of k_u_ values. The 95% confidence interval is shown in red as [157.5, 202.5] [s].

**Fig 3.**
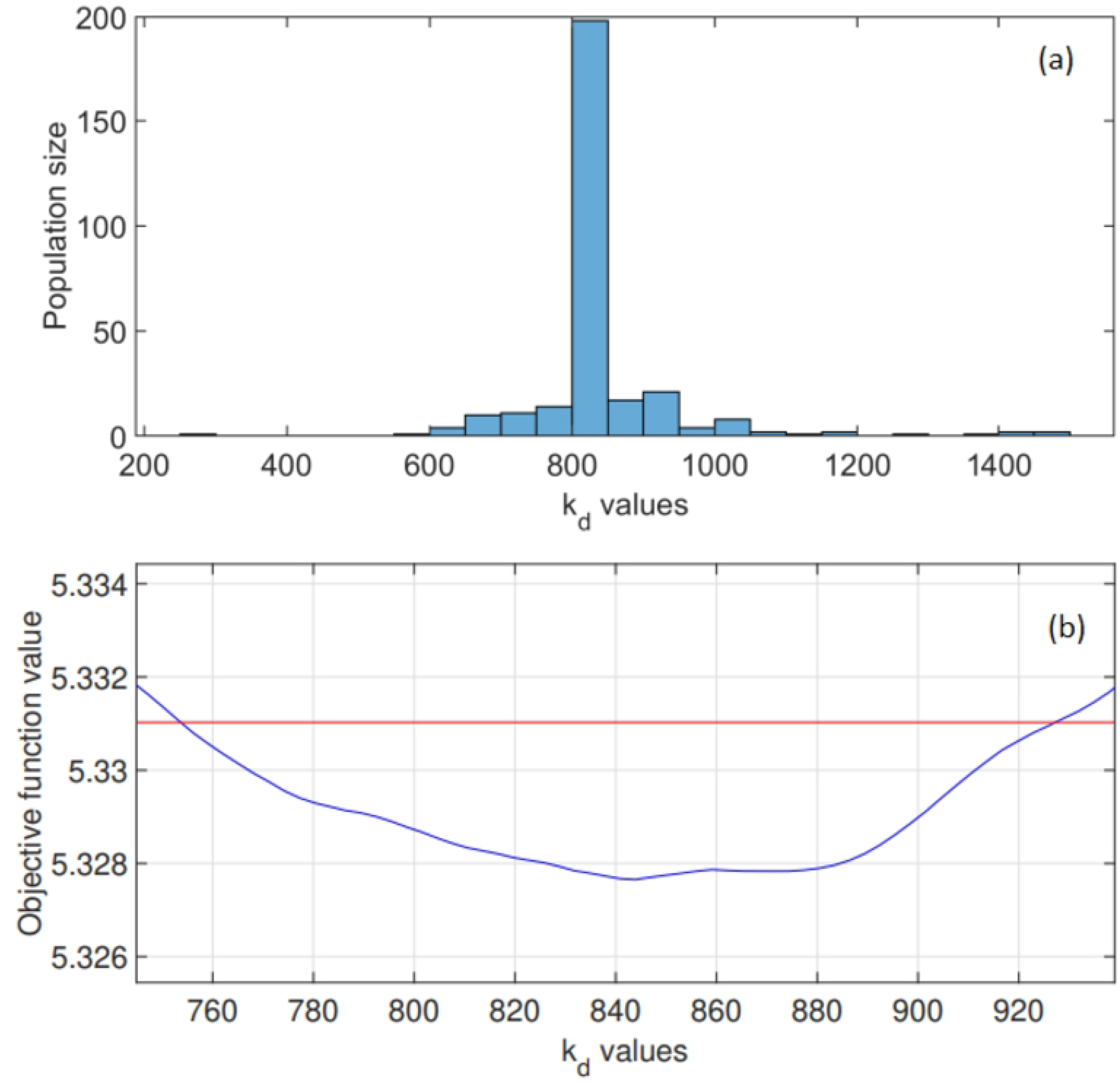
Global optimisation results for the stomatal parameter k_d_ in expression (16). (a) The range of potential k_d_ values considered is shown along with the optimised k_d_ distribution. The genetic algorithm converged to a parameter on the interval [800,850], with the optimised k_d_=830.3 [s]. (b) Objective function values: computed profile likelihood for a range of k_d_ values. The 95% confidence interval is shown in red as [735.6, 927.5] [s].

**Fig 4.**
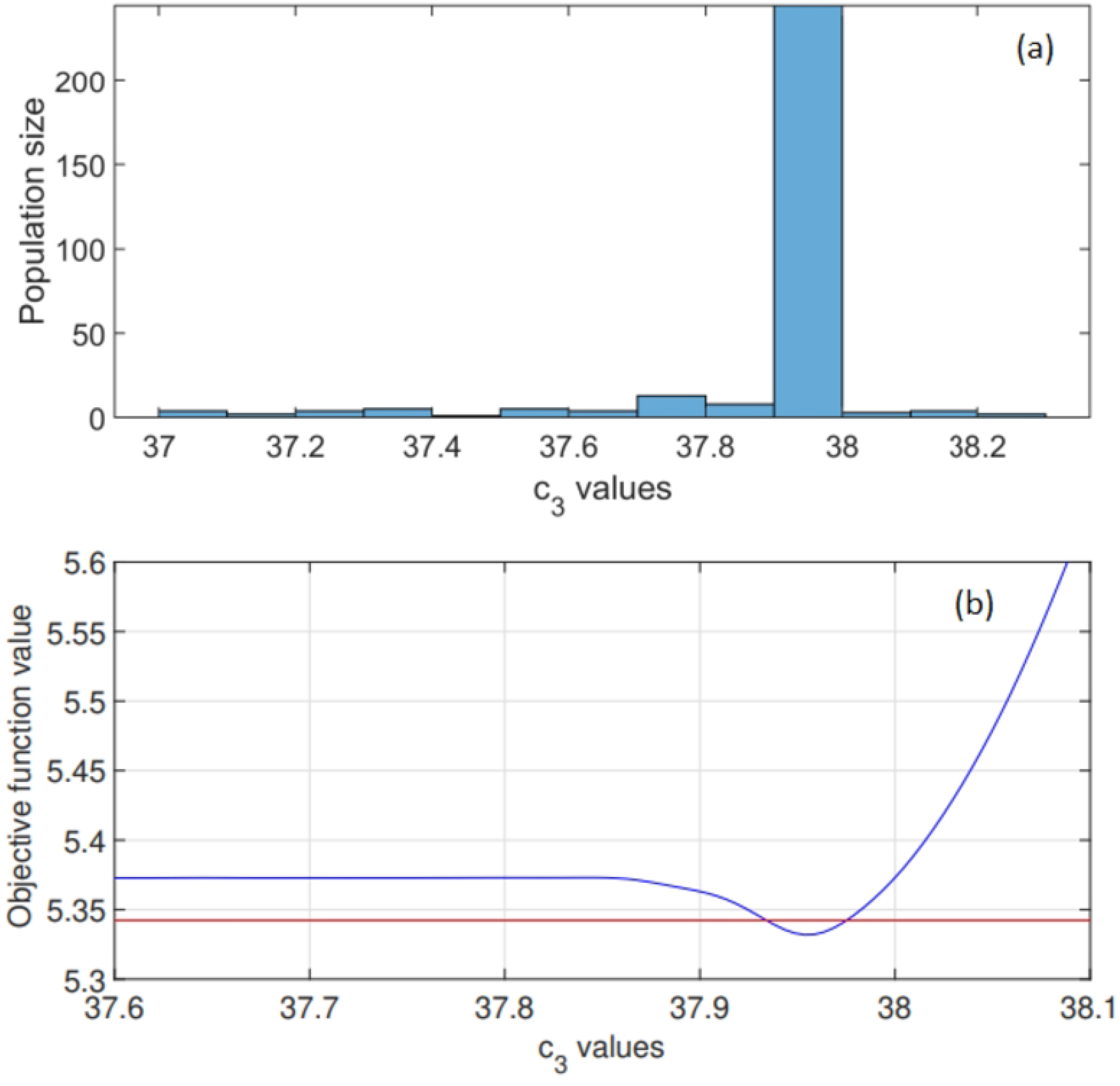
Global optimisation results for parameter c_3_ in expression (10) used in (18). (a) The range of potential c_3_ values considered is shown along with the optimised c_3_ distribution. The genetic algorithm converged to an optimal parameter value on the interval [37.9,38] as c_3_=37.96. (b) Objective function values: computed profile likelihood for a narrow range of c_3_ values. The 95% confidence interval is shown in red as [37.95, 37.98] [s].

Notice that the derived time constants k_u_=179.4 [s] and k_d_=830.3 [s] suggest that for a leaf under natural fluctuating irradiance, the overall stomatal dynamics related to an increase in irradiance is faster than those associated with a decrease in irradiance.

The optimised c_3_ value 37.96 (see Fig.(4)) increases the value of W_c_ (defined in expression (4)) compared to this function’s value computed using the commonly used c_3_=38.28 [27]. All parameter values in (15) and (16) are given in Table (2) with parameter values in (18) given in Table (3).

**Table 2.** Values of the model parameters in expressions (15) and (16). Parameters of the steady-state target function G(I(t),c_a_(t)) (indicated in blue), computed under c_a_=400ppm and c_a_=800ppm respectively, are also included. Estimated unknown system parameters are indicated in red with accompanying confidence intervals. Steady-state parameters estimated *a priori* are indicated in green.

| Parameter | Description | Value |
| --- | --- | --- |
| $k_u$ [s] | Time constant for increases in $g_{tc}$ | 179.4 [157.5,202.5] |
| $k_d$ [s] | Time constant for decreases in $g_{tc}$ | 830.3 [735.6,927.5] |
| $G(I(t), c_a(t))$ [$\frac{mol}{m^2 s}$] | Steady-state target function for stomatal conductance | [S1 Supporting Information] |
| $g_{min,400}$ [$\frac{mol}{m^2 s}$] | Minimum stomatal conductance under $c_a = 400\text{ppm}$ | 0.1 |
| $g_{max,400}$ [$\frac{mol}{m^2 s}$] | Maximum stomatal conductance under $c_a = 400\text{ppm}$ | 0.325 |
| $\alpha_{400}$ [$\frac{mol}{\mu mol}$] | Slope of relation between steady-state $G$ and $I$ under $c_a = 400\text{ppm}$ | 0.00033 |
| $g_{min,800}$ [$\frac{mol}{m^2 s}$] | Minimum stomatal conductance under $c_a = 800\text{ppm}$ | 0.165 |
| $g_{max,800}$ [$\frac{mol}{m^2 s}$] | Maximum stomatal conductance under $c_a = 800\text{ppm}$ | 0.29 |
| $\alpha_{800}$ [$\frac{mol}{\mu mol}$] | Slope of relation between steady-state $G$ and $I$ under $c_a = 800\text{ppm}$ | 0.00017 |

**Table 3.**
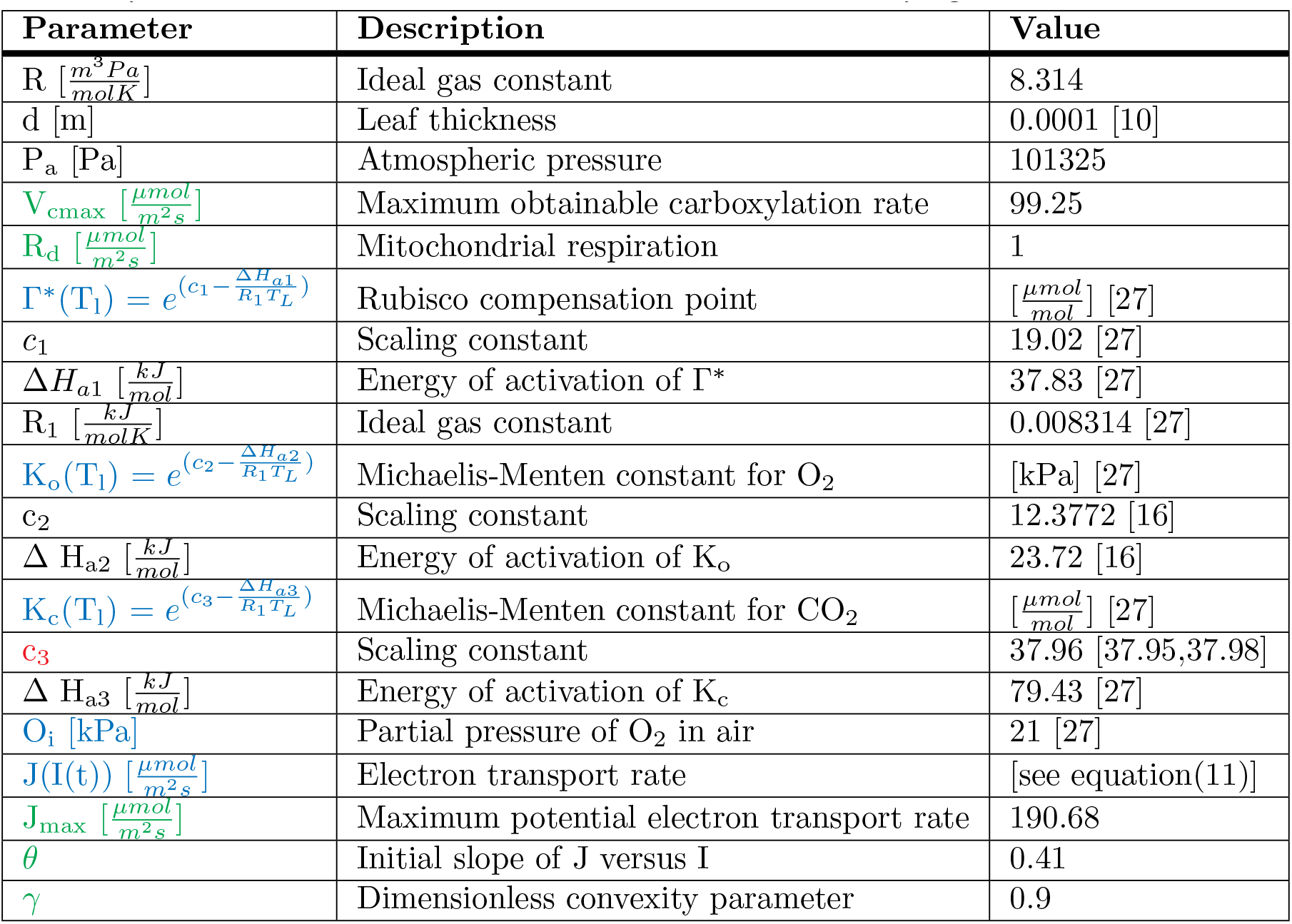
Values of the model parameters in expression (18). *A priori* estimated parameters of function J(I(t)) and the respective steady-state parameters V_cmax_ and R_d_ are shown in green. The respective functions are indicated in blue and the estimated unknown system parameter is indicated in red with its accompanying confidence interval.

**Table 4.** Greenhouse data measured under natural fluctuating irradiance. Summary of RMSEs computed to assess the accuracy of A_n_. Here 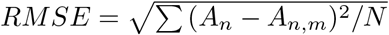, with the measured output denoted by A_n,m_. Results comparing the use of the optimised parameter c_3_=37.96 to the original c_3_=38.28 [27] are shown. Experiments were conducted under 2 sets of ambient CO_2_ concentrations, 400ppm and 800ppm respectively. R^2^ values are given in [ ].

| Data set ($c_a=400\text{ppm}$ RH=75%) | RMSE with $c_3=37.96$ | RMSE with $c_3=38.28$ |
| --- | --- | --- |
| Estimation set (8 Sept 2021) (Fig. 5) | 0.747 [0.98] | 2.082 [0.85] |
| Validation set 1 (3 Sept 2021) | 0.661 [0.98] | 1.54 [0.88] |
| Validation set 2 (6 Sept 2021) (Fig. 7) | 0.659 [0.99] | 1.3 [0.95] |
| Validation set 3 (23 Sept 2021) | 0.998 [0.9] | 0.936 [0.91] |
| Validation set 4 (24 Sept 2021) | 1.164 [0.91] | 1.13 [0.92] |
| Data set ($c_a=800\text{ppm}$ RH=75%) | | |
| Validation set 5 (7 Sept 2021) (Fig. 8) | 0.89 [0.97] | 0.89 [0.97] |

### 4.2 Photosynthesis under naturally fluctuating irradiance: parameter estimation

We parameterise the model using data measured on 8 September 2021 under natural fluctuating light as this contains rich information pertaining to fluctuations between W_c_ and W_j_ limitation (refer to Fig.(5.c)).

**Fig 5.**
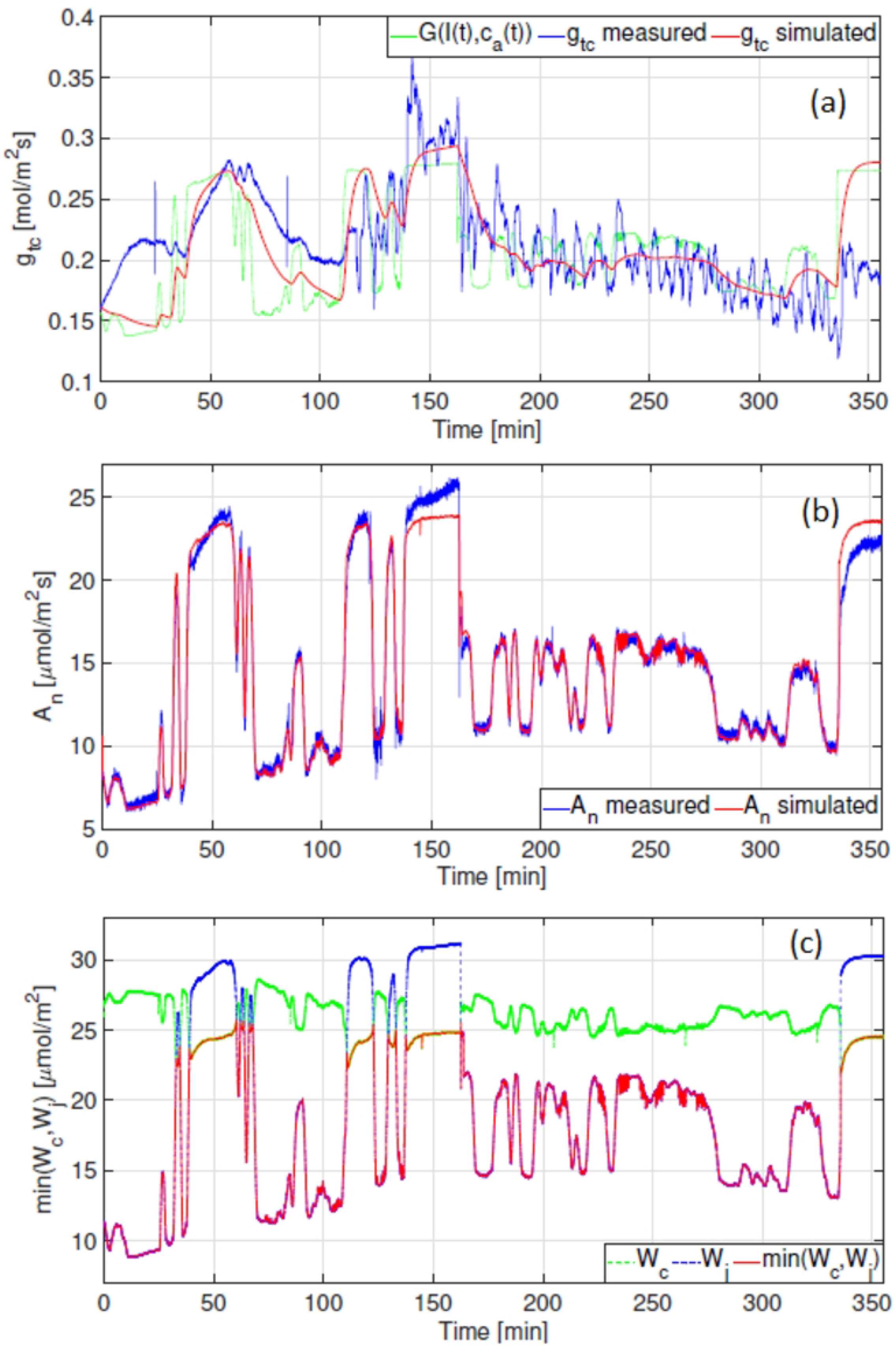
Results obtained after parameter estimation. (a) Total stomatal conductance to CO_2_ diffusion [*molm*^−2^*s*^−1^]. (b) Net photosynthetic rate [*μmolm*^−2^*s*^−1^] (c) min(W_*c*_,W_*j*_) [*μmolm*^−2^*s*^−1^], indicates whether photosynthesis is limited by the carboxylation or electron transport rate.

Predictions for g_tc_ are shown in red in Fig.(5.a). A 9% increase in predicted accuracy (from the objective function in (19)) was obtained by modelling g_tc_ with an asymmetric as opposed to a symmetric response to irradiance (see S1 Supporting Information).

The dynamic relationship between the respective W_c_ and W_j_ limitations is shown in red in Fig.(5.c). Results indicate that Rubisco limited photosynthesis (W_c_) coincides with elevated levels of natural irradiance seen Fig.(1.a). The model predicts that photosynthesis is limited by the electron transport rate (W_j_) for the majority of the day, thus for irradiance levels that remain below 400 [*μmolm*^−2^*s*^−1^]. The correlation between A_n_ and the rate of electron transport J(I(t)) is shown in Fig.(6).

**Fig 6.**
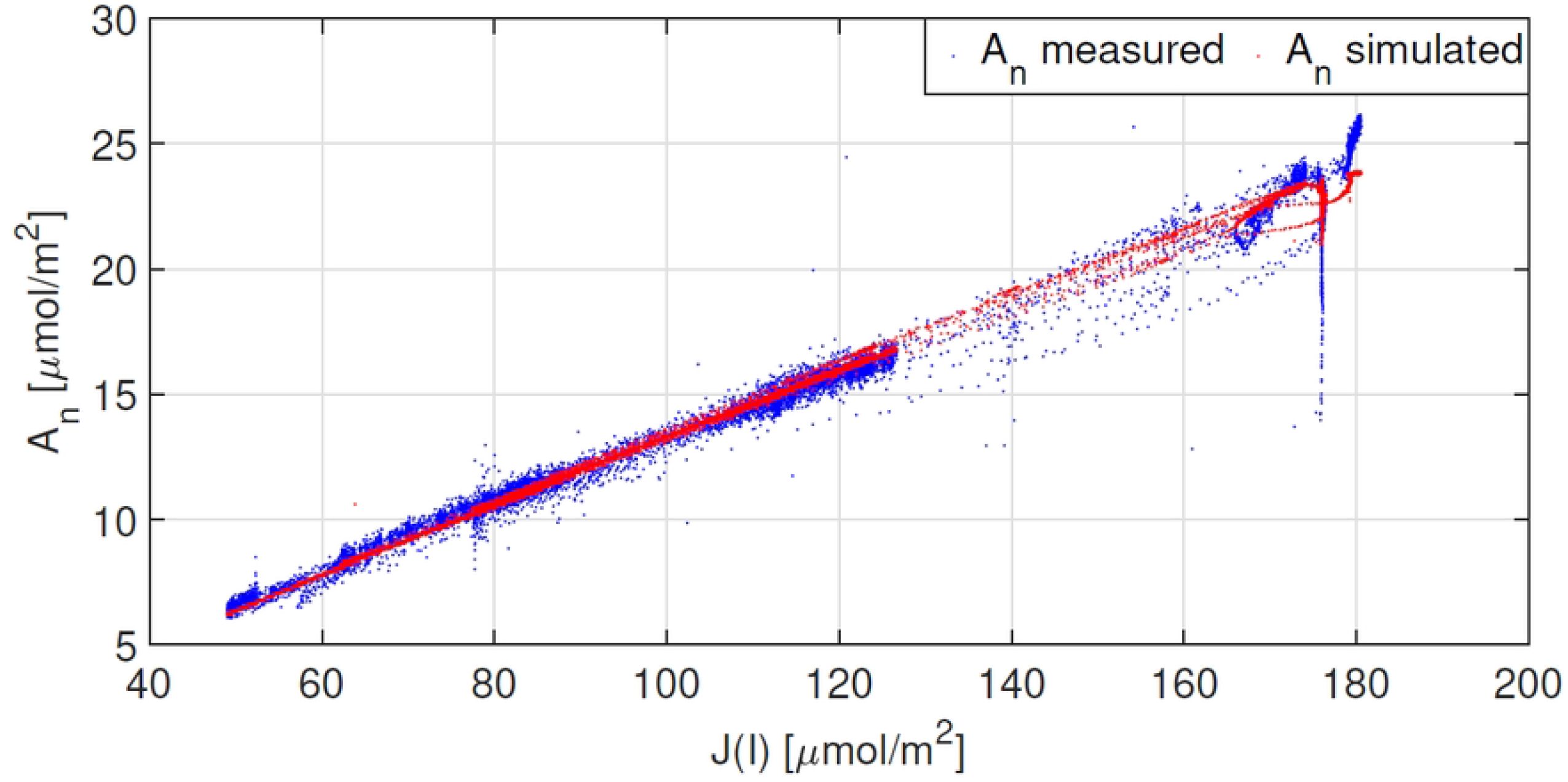
Results obtained after parameter estimation. The correlation between A_n_ and the electron transport rate (J(I(t)) is shown for both measured and simulated data sets.

### 4.3 Photosynthesis under natural fluctuating irradiance: model validation

We proceed by using the parameter values obtained in section (4.2) (given in Tables (2) and (3)) during model validation. The outcome is summarised in Table (4).

The results obtained for measurements taken on 6 Sept 2021 under an ambient CO_2_ concentration of 400ppm are shown in Fig.(7), whilst the results for measurements conducted on 7 Sept 2021 under a CO_2_ concentration of 800ppm are shown in Fig.(8). Both Figs.(7.b) and (8.b) show good agreement between measured and modelled A_n_. Simulations shown in Fig.(8.c) suggest that when c_a_ is 800ppm, photosynthesis is solely limited by the electron transport rate. Furthermore, one observes similarities between the shapes of the modelled W_j_ and A_n_. Given that W_jc_ is computed using J(I(t)) from expression (11), this suggests that for tomato under the conditions reported here, it may be sufficient to use electron transport rates, inferred from steady-state data, in a dynamic setting. The correlation between A_n_ and J(I(t)) is shown in Fig.(9).

**Fig 7.**
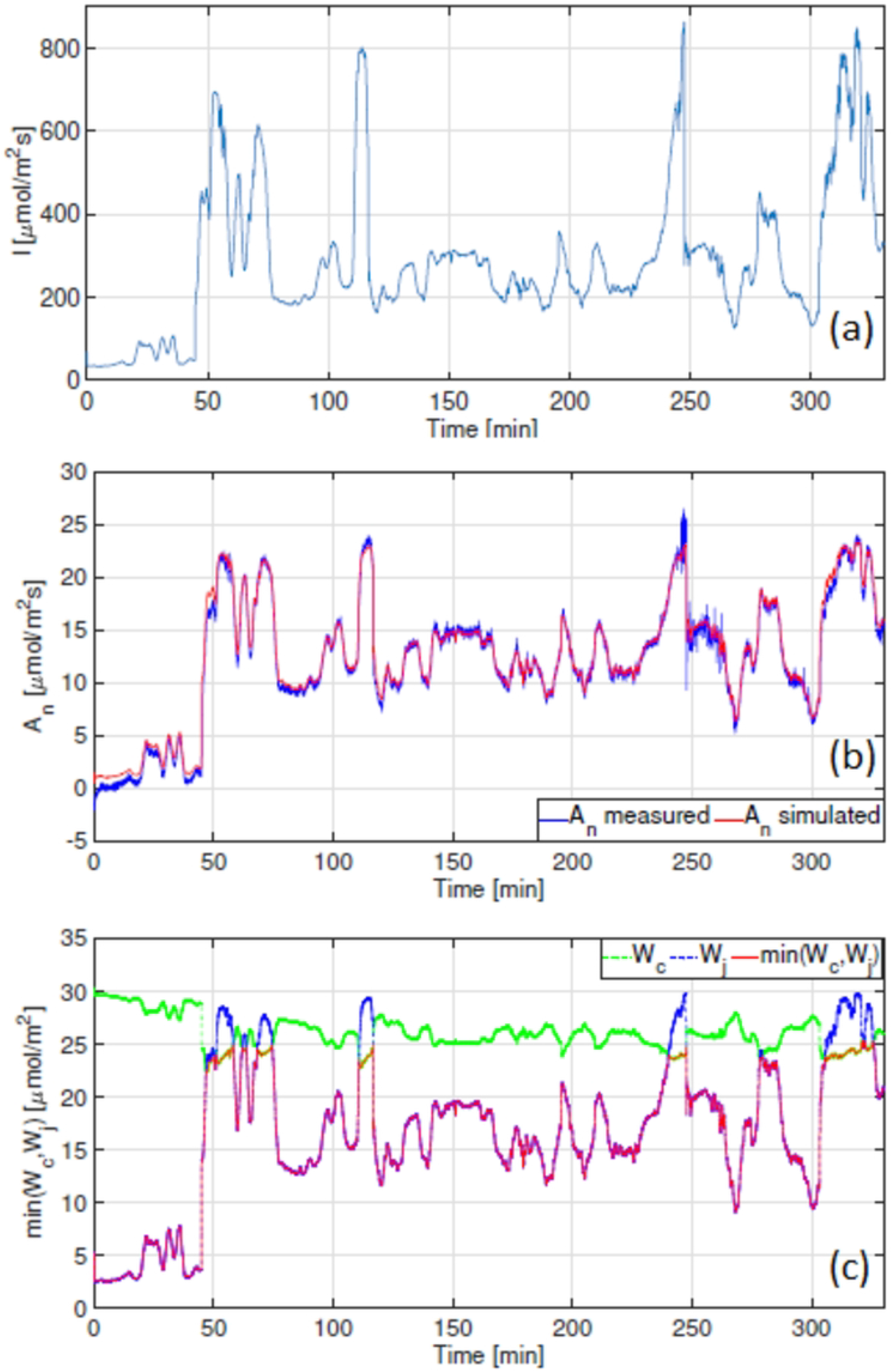
Model validation results obtained for measurements taken on 6 Sept 2021 with c_a_=400ppm. (a) Measured irradiance [*μmolm*^−2^*s*^−1^]. (b) Net photosynthetic rate [*μmolm*^−2^*s*^−1^] (c) min(W_*c*_,W_*j*_) [*μmolm*^−2^*s*^−1^], indicates whether photosynthesis is limited by the carboxylation or electron transport rate.

**Fig 8.**
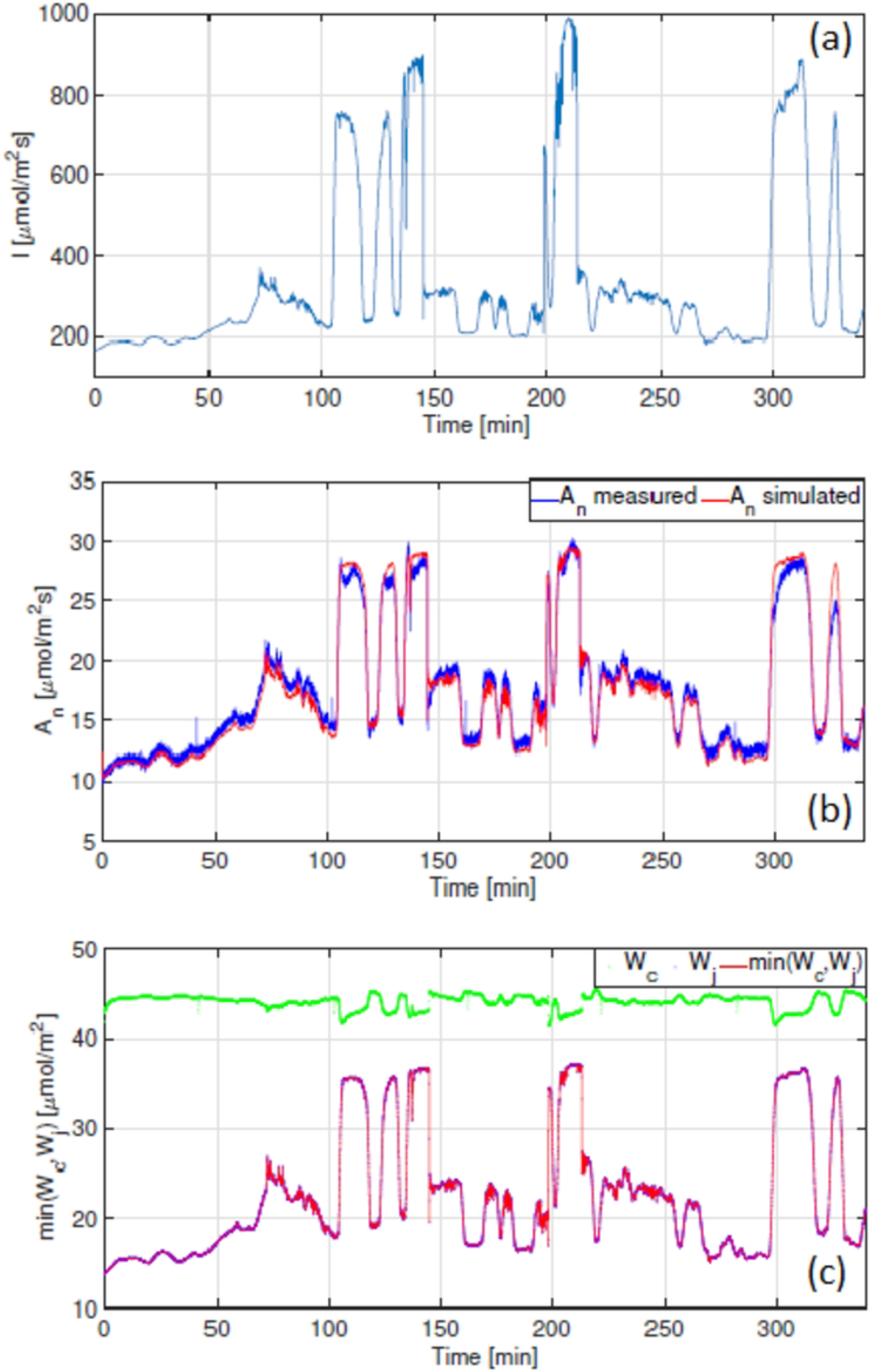
Model validation results obtained for measurements taken on 7 Sept 2021 with c_a_=800ppm. (a) Measured irradiance [*μmolm*^−2^*s*^−1^]. (b) Net photosynthetic rate [*μmolm*^−2^*s*^−1^] (c) min(W_*c*_,W_*j*_) [*μmolm*^−2^*s*^−1^], indicates whether photosynthesis is limited by the carboxylation or electron transport rate.

**Fig 9.**
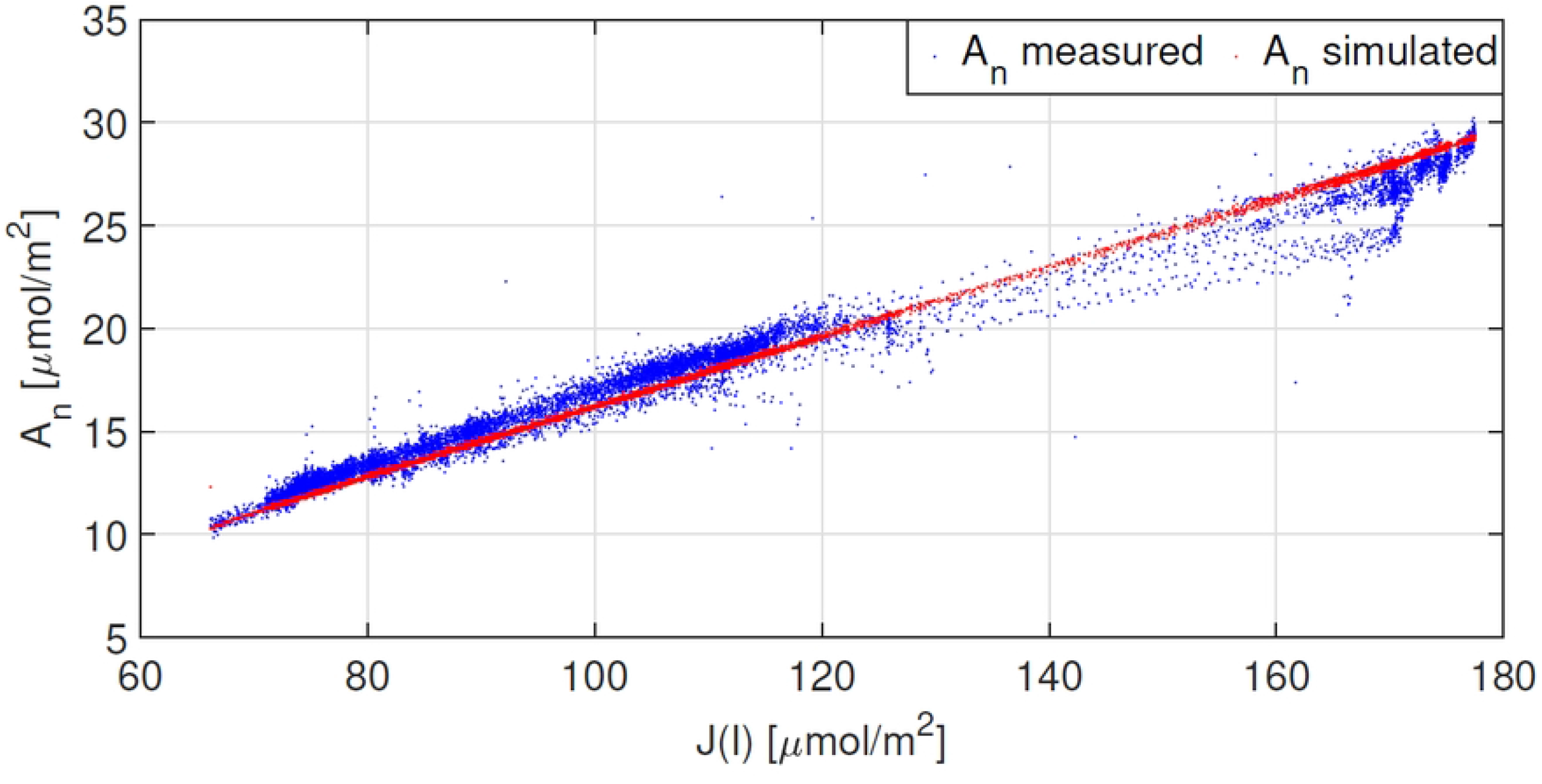
Results obtained for the validation data set measured on 7 Sept 2021 under elevated c_a_ = 800ppm. The correlation between A_n_ and the electron transport rate (J(I(t)) is shown for both measured and simulated data sets.

## 5 Discussion

It is highly unlikely that A_n_ is in steady-state under natural fluctuating irradiance conditions and so observing natural dynamic as opposed to step-change responses to light is useful in aiding our understanding of this key photosynthetic property. However, nonsteady-state photosynthesis is often overlooked, with kinetic measurements of A_*n*_ reported less due to the complexity associated with measuring and analysing them [28].

We set out to develop a fit-for-purpose dynamic photosynthesis model. The model is both calibrated and validated using measurements taken under naturally fluctuating greenhouse conditions. Sufficiently accurate A_n_ predictions in section (4) suggest that the model (given in expressions (15)-(17)) can potentially be used in greenhouse lighting control applications.

### Model

Our model comprises 2 ODEs, predicting the total stomatal conductance to CO_2_ (g_tc_) diffusion and the CO_2_ concentration inside a leaf (c_i_). These predictions are required to compute the net photosynthetic rate (A_n_) using Fick’s law of diffusion (expression (14)). Our results show that satisfactory fit for purpose A_n_ values can be obtained by merely predicting the elements that comprise Fick’s law of diffusion.

The dynamic binding of CO_2_ inside a leaf is modelled using the FvCB model that has been adapted the predict steady-state J values at different light intensities (see section (1.3)). Here, we used this application in a dynamic setting, defining the dynamic electron transport rate function, J(I(t)). It does not include the process of how J gradually increases after light increase.

Given that parameter V_cmax_ is estimated from *a priori* A/c_i_ measurements (see S1 Supporting Information for details), we chose to optimise a parameter used to describe the temperature dependence of the Michaelis-Menten constant for CO_2_, c_3_, to increase the accuracy of our A_n_ predictions (see Table (4)).

The model is unique given its small size and the fact that it only comprises 3 unknown system parameters. We opt not to predict any detailed molecule concentrations such as RuBP (see [19] for example), and by assuming that the leaf is homogeneous, we do not include an additional equation that accounts for mesophyllic conductance (see [11] for example).

Furthermore, for V_cmax_ as defined in the FvCB model [7], we too assume is that Rubisco is activated [29]. Predictions for A_n_ reported for tomato suggest that adequate A_n_ values can be obtained under this stringent model assumption.

Finally, it is important to realise that the implementation of a system which contains the FvCB model necessitates the *a priori* calculation of a multitude of species specific steady-state parameters. This requires repeated experimental measurements and some background knowledge on how to interpret and assimilate data.

### Results

The first interesting point that emerges from using rapid fluctuating measurements to calibrate a model is the inferred time constants pertaining to the total stomatal conductance to CO_2_ diffusion. Notice from Table (2) that k_u_, associated with an increase in irradiance is faster than k_d_. When inferring these parameter values from data measured after a single step change in irradiance, k_u_ is slower than k_d_.

We highlighted the correlation between A_n_ and the electron transport rate in section (4) (see Figs. (6) and (9)). We know that the formation of ATP and NADPH molecules are dependent on the rate of electron transport and that this is light dependent. Accordingly, we observe a strong linear relationship between between A_n_ and J under low irradiance levels. This indicates that under such conditions, photosynthesis is W_*j*_ limited. Accurate results shown in Figs.(7) and (8.b) in particular, suggest that: 1) it is sufficient to use an electron transport rate function, at least for our greenhouse-grown tomato leaves, calibrated using steady-state values at different irradiance levels, in a dynamic setting, and 2) the calibration of this function i.e. the parameter values J_max_, *θ* and *γ*, is critically important.

## 6 Conclusions

The importance of monitoring nonsteady-state responses to natural fluctuations in irradiance for improving crop photosynthesis has gained substantial support in recent years [30]. Our aim was to develop a small fit-for-purpose dynamic photosynthesis model that can be used in supplemental lighting control applications in greenhouses. We set out to build a model that accurately predicts the net photosynthetic rate (A_n_) by taking plant physiology into account, and both calibrated and validated our model using nonsteady-state data measured under rapid fluctuating light conditions.

Four main points have emerged from our analysis:

1. We corroborated the added value of accounting for differences in stomatal responses to both increasing and decreasing fluctuations in irradiance. We observed a 9% increase in model accuracy when using 2 different time constants to describe the total stomatal conductance to CO_2_ (refer to S1 Supporting Information). In contrast, we observed no significant increase in the predicted accuracy of A_*n*_ when modelling the steady-state target function of the total stomatal conductance to CO_2_, denoted by G, as a function of both irradiance and ambient CO_2_ concentration (refer to S1 Supporting Information).
2. We showed that incorporating the FvCB equations into a dynamic model is sufficient for obtaining accurate fit for purpose A_n_ predictions under rapid natural light fluctuations. In particular, we found that the *a priori* parameterisation of the steady-state electron transport rate with respect to different irradiance levels (J(I(t)) is very effective in capturing the dynamics of photosynthesis in tomato leaves.
3. We showed the added value of optimising a parameter in one of the Michaelis-Menten constants in the FvCB model. Here, we opted to adjust parameter c_3_ related to K_n_. Its value is often used from literature, despite being calibrated for different plant species. This cautions us when we are simply using parameter values from literature derived for different plant species.
4. We showed that satisfactory photosynthesis results can be obtained even when a model does not account for complex biological factors such as enzymatic inhibition, liquid-gas interactions or chloroplast movement.

## Nomenclature

*Symbols*

A_n_: Net photosynthetic rate [*μmolm*^−2^*s*^−1^]
g_tc_: Total stomatal conductance to CO_2_ diffusion [*molm*^−2^*s*^−1^]
g_m_: Mesophyllic conductance [*molm*^−2^*s*^−1^]
c_i_: CO_2_ concentration inside a leaf [*μmolm*^−2^*s*^−1^]
c_a_: Ambient CO_2_ concentration [*μmolm*^−2^*s*^−1^]
c_c_: CO_2_ concentration in the chloroplast stroma [*μmolm*^−2^*s*^−1^]
I: Irradiance [*μmolm*^−2^*s*^−1^]
T_l_: Leaf temperature [K]
k_u_: Time constant for increases in g_tc_ [s]
k_d_: Time constant for decreases in g_tc_ [s]
R: Ideal gas constant [*m*^3^*Pa*^1^*mol*^−1^*K*^−1^]
d: Leaf diameter [m]
P_a_: Atmospheric pressure [Pa]
V_cmax_: Maximum obtainable carboxylation rate [*μmolm*^−2^*s*^−1^]
K_c_: Michaelis-Menten constant for CO_2_ [*μmolm*^−2^*s*^−1^]
K_o_: Michaelis-Menten constant for O_2_ [kPa]
O_i_: Partial pressure of O_2_ in air [kPa]
Γ*: The Rubisco compensation point [*μmolm*^−2^*s*^−1^]
J: Steady-state electron transport rate [*μmolm*^−2^*s*^−1^]
G: Steady-state total stomatal conductance to CO_2_ diffusion [*molm*^−2^*s*^−1^]
W_c_: Rubisco limited carboxylation rate [*μmolm*^−2^*s*^−1^]
W_j_: Electron transport rate limited carboxylation rate [*μmolm*^−2^*s*^−1^]
R_d_: Mitochondrial respiration which includes CO_2_ release in light other than photo-respiration [*μmolm*^−2^*s*^−1^]
RH: Relative humidity [%]

## Supporting information

**S1 Supporting Information. A small dynamic leaf-level model predicting photosynthesis in greenhouse tomatoes**.

